# Uncultivated DPANN archaea are ubiquitous inhabitants of global oxygen deficient zones with diverse metabolic potential

**DOI:** 10.1101/2023.10.30.564641

**Authors:** Irene H. Zhang, Benedict Borer, Rui Zhao, Steven Wilbert, Dianne K. Newman, Andrew R. Babbin

## Abstract

Archaea belonging to the DPANN superphylum have been found within an expanding number of environments and perform a variety of biogeochemical roles, including contributing to carbon, sulfur, and nitrogen cycling. Generally characterized by ultrasmall cell sizes and reduced genomes, DPANN archaea may form mutualistic, commensal, or parasitic interactions with various archaeal and bacterial hosts, influencing the ecology and functioning of microbial communities. While DPANN archaea reportedly comprise 15–26% of the archaeal community within marine oxygen deficient zone (ODZ) water columns, little is known about their metabolic capabilities in these ecosystems. We report 33 novel metagenome-assembled genomes belonging to DPANN phyla Nanoarchaeota, Pacearchaeota, Woesarchaeota, Undinarchaeota, Iainarchaeota, and SpSt-1190 from pelagic ODZs in the Eastern Tropical North Pacific and Arabian Sea. We find these archaea to be permanent, stable residents of all 3 major ODZs only within anoxic depths, comprising up to 1% of the total microbial community and up to 25–50% of archaea. ODZ DPANN appear capable of diverse metabolic functions, including fermentation, organic carbon scavenging, and the cycling of sulfur, hydrogen, and methane. Within a majority of ODZ DPANN, we identify a gene homologous to nitrous oxide reductase. Modeling analyses indicate the feasibility of a nitrous oxide reduction metabolism for host-attached symbionts, and the small genome sizes and reduced metabolic capabilities of most DPANN MAGs suggest host-associated lifestyles within ODZs.

**Importance:** Archaea from the DPANN superphylum have diverse metabolic capabilities and participate in multiple biogeochemical cycles. While metagenomics and enrichments have revealed that many DPANN are characterized by ultrasmall genomes, few biosynthetic genes, and episymbiotic lifestyles, much remains unknown about their biology. We report 33 new DPANN metagenome-assembled genomes originating from the 3 global marine oxygen deficient zones (ODZs), the first from these regions. We survey DPANN abundance and distribution within the ODZ water column, investigate their biosynthetic capabilities, and report potential roles in the cycling of organic carbon, methane, and nitrogen. We test the hypothesis that nitrous oxide reductases found within several ODZ DPANN genomes may enable ultrasmall episymbionts to serve as nitrous oxide consumers when attached to a host nitrous oxide producer. Our results indicate DPANN archaea as ubiquitous residents within the anoxic core of ODZs with the potential to produce or consume key compounds.

## Introduction

In recent years, metagenomics has enabled the discovery of several prokaryotic superphyla lacking pure culture representatives (1–3). One of these novel groups is the DPANN archaea, named after the first members of the expanding superphylum (Diapherotrites, Parvarchaeota, Aenigmarchaeota, Nanoarchaeota and Nanohaloarchaeota) which has come to include at least ten putative phyla (4, 5). The DPANN archaea are characterized by ultrasmall cell sizes (∼0.1–1.5 µm), reduced genomes (∼1.5 Mb), and limited metabolic capacities (6). These features, along with several enrichments and visualizations of DPANN archaeal-host associations (7–9), suggest a symbiotic or commensal lifestyle of DPANN archaea with diverse microbial hosts. If DPANN indeed exist in partnership with others, this would explain why they have been challenging to cultivate in isolation.

Since their discovery, DPANN archaea have been found in a variety of diverse environments, including hydrothermal vents (10), freshwater and hypersaline lakes (11, 12), groundwater (13, 14), terrestrial hot springs (15), marine sediments and water columns (10, 16, 17), and the Black Sea (18). Archaea writ large play crucial roles in global biogeochemical cycles, such as in ammonia oxidation (19), methane cycling (20), and organic carbon scavenging (21), and DPANN archaea have been found to possess genes for sulfur cycling and organic substrate degradation (13, 16). Additionally, DPANN archaea in anoxic environments may form consortia with methanogens and contribute to anaerobic carbon cycling (22). However, despite their widespread abundance, distribution, and diversity (accounting for about half of all archaeal diversity (6)), the ecological and biogeochemical roles of DPANN archaea are not fully understood. Culture-independent techniques have only begun to unravel the importance of these previously-overlooked microorganisms within their expanding list of habitats.

Amplicon surveys have detected the presence of DPANN archaea within both sediments beneath oxygen deficient zones (ODZs) (23) and the ODZ water column itself (24). The three major oceanic ODZs are located in the eastern tropical North Pacific (ETNP), the eastern tropical South Pacific (ETSP), and the Arabian Sea. Oxygen profiles in these regions display rapid decreases from surface saturation to below the detection limit of trace oxygen sensors (<10 nmol L^-1^) between 50–100 m depth, a region termed the oxycline (25, 26). Oxygen concentrations then remain below detection and with no vertical gradient for approximately 200–800 m (27), although the ODZ thickness varies greatly across each basin (28–30). Due to these unique features, ODZ water columns contain multiple biogeochemical gradients that support diverse microbial assemblages performing nitrogen, carbon, and sulfur cycling (31). In particular, these regions disproportionately contribute to marine nitrogen cycling, accounting for about 30% of marine fixed nitrogen loss despite containing only 0.1–0.2% of oceanic volume (32, 33).

ODZs are characterized by prevalent denitrification, *i.e.* the microbially-mediated stepwise reduction of nitrate to dinitrogen gas. This anaerobic respiratory metabolism occurs via reductases encoded by a suite of widely distributed genes (34). The last step of denitrification, the reduction of N_2_O to N_2_, is catalyzed by nitrous oxide reductase encoded by *nos.* Two clades of the *nos* catalytic subunit *nosZ* have been found, a typical clade I *nosZ* associated with complete denitrifiers defined by an N-terminal twin-arginine translocation (TAT) motif, and an atypical clade II *nosZ* associated with partial denitrifiers defined by an N-terminal Sec-type motif (35). Both variants contain conserved copper-binding sites Cu_A_ and Cu_Z_, although Cu_Z_ sites of clade II *nosZ* homologs exhibit greater variability and less conservation (36). Recent studies reveal clade II *nosZ* predominates within ODZs, occurs within diverse marine taxa including archaea, and may be associated with low oxygen and enhanced N_2_O affinity (36). Because N_2_O depletes ozone and is a potent greenhouse gas, organisms with atypical *nosZ* variants, including archaea, merit interest as potential N_2_O sinks.

Increasing attention has been focused on ODZ archaeal communities (37–39), such as members of Thermoproteota (including former Marine Group I Thaumarchaeota) and Thermoplasmatota (including former Marine Group II archaea) (40, 41). However, little is known about ODZ DPANN archaea, despite reports that they may comprise up to 15–26% of total archaeal reads in these regions (24). Challenges in cultivation of these environmental microbes limit our understanding of the metabolic capabilities of clades such as DPANN that lack cultured representatives. Accordingly, the contribution of DPANN archaea in ODZ microbial assemblages and biogeochemical cycling, as well as the abundance, distribution, metabolism, ecology, and phylogeny of these archaea remain open questions. Using genome-resolved metagenomics, we recover 33 genomes belonging to DPANN phyla Nanoarchaeota, Pacearchaeota, Woesarchaeota, Undinarchaeota, and Iainarchaeota from the ETNP and Arabian Sea ODZs. We characterize the metabolic capabilities of these archaea, place them within the existing phylogeny of known DPANN, and determine their relative abundances and distributions within and across global ODZs. Our results demonstrate that DPANN are a ubiquitous portion of the microbial community within ODZs and comprise several lineages with diverse metabolic potential.

## Materials and Methods

### Sample collection, sequencing, metagenome assembly, and binning

Sampling and sequencing methods for public ETNP metagenomes are described in Fuchsman *et al.* 2017 (39), Glass *et al.* 2015 (42), and Tsementzi *et al.* 2016 (43). Sampling and sequencing methods for public ETSP metagenomes are described in Stewart *et al.* 2012 (44) and Ganesh *et al.* 2014 (45). Raw reads per metagenome were retrieved from the Sequence Read Archive (SRA) using the following NCBI BioProject IDs: PRJNA350692 (Fuchsman ETNP metagenomes), PRJNA254808 (Glass ETNP metagenomes), PRJNA323946 (Tsementzi ETNP metagenomes), PRJNA68419 (Stewart ETSP metagenomes), and PRJNA217777 (Ganesh ETSP metagenomes. Sampling locations for each metagenome were visualized using Python 3.7.12 and the cartopy package. These were plotted against global oxygen concentrations from 300 m below sea surface from Ocean Data Atlas 2018 (Figure 1A).

**Figure 1:**
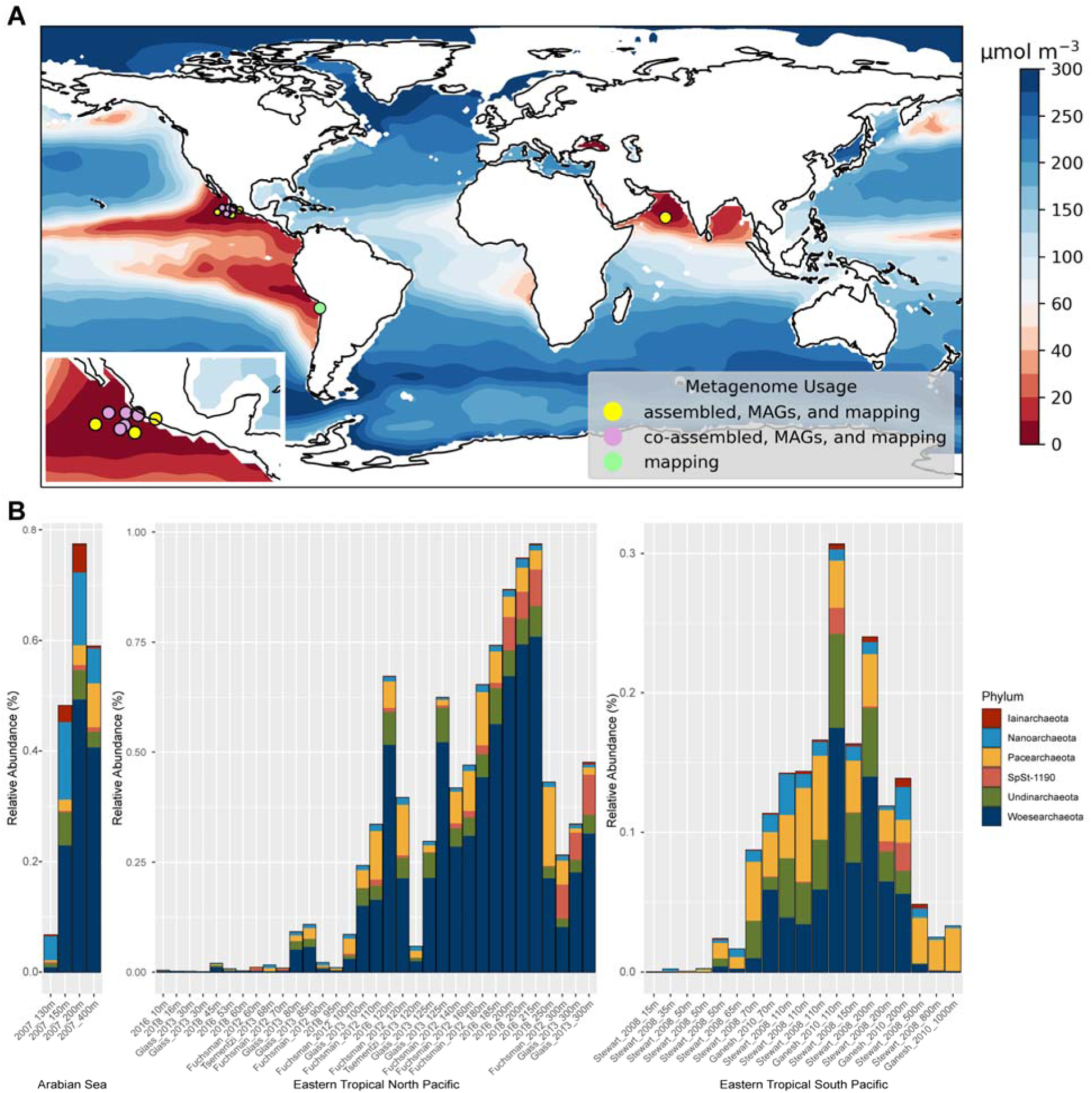
**A)** Locations of metagenomes from ETNP, ETSP, and Arabian Sea used for metagenome assembly, MAG binning, and relative abundance mapping **B)** Relative abundances of DPANN MAGs across metagenome samples, color-coded by phylum-level taxonomy.

Trimming of raw reads, metagenome assembly, and binning methods are described elsewhere (46). Metagenome-assembled genomes (MAGs) were defined as bins with completion >50% and contamination <10% according to CheckM (47), although these statistics based on single-copy genes may underestimate the true completeness of DPANN archaea MAGs due to their limited genome sizes. Taxonomy was assigned to all MAGs using GTDB-tk v1.7.0 with the classify_wf workflow (48). 33 MAGs belonging to DPANN phyla were annotated with PROKKA v1.14.6 (49) against the HAMAP (50) and Pfam databases (51) using the --kingdom Archaea flag. The full set of ODZ MAGs, as well as all DPANN MAGs generated in this study, were deposited under NCBI BioProject ID PRJNA955304.

Published DPANN MAGs and genomes were manually downloaded from the Joint Genome Institute (JGI), while published DPANN from the National Center for Biotechnology Information (NCBI_ were downloaded using the EntrezDirect utility. These DPANN MAGs and genomes were assessed for completeness and contamination with CheckM v1.0.12 (47), and detailed taxonomy was determined with GTDB-tk v1.7.0 (48). MAGs and genomes below 50% completion and above 10% contamination, along with those that did not taxonomically classify within DPANN phyla, were pruned and remaining genomes were dereplicated with dRep v3.2.2 (52) with the -sa 0.99 flag to remove redundant genomes.

TARA Oceans MAGs were retrieved from Delmont et al. 2018 (53). To determine if DPANN MAGs were present within the TARA Oceans collection, we reclassified the 957 non-redundant TARA Oceans MAGs with GTDB-tk v1.7.0 (48). These taxonomies were then searched for the presence of any DPANN phyla.

For dereplicated ODZ DPANN MAGs, coverage mapping was performed with CoverM using the flags minimap2-sr --min-read-aligned-percent 50 --min-read-percent-identity 0.95 -- min-covered-fraction 0 (https://github.com/wwood/CoverM). Relative abundances of dereplicated MAGs resulting from CoverM mapping were visualized using R v4.1.3 and the packages phyloseq, ggplot2, and dplyr.

### Gene searching, metabolic analysis, and tree building

Unique published DPANN and all ODZ DPANN MAGs were queried for 76 archaea-specific single copy genes, which were aligned using GtoTree v1.6.31 with the -H Archaea -G 0.25 flags (54). We created a phylogenetic tree based on the output archaeal single copy gene alignment with IQ-Tree v1.6.12 (55) using the WAG+R6 model and 1000 ultrafast bootstraps (56).

To determine metabolic capabilities of ODZ DPANN, we used Anvi’o 7.1 (57). Briefly, for each DPANN MAG we generated a contigs database with anvi-gen-contigs-database. For metabolic predictions, we ran anvi-run-kegg-kofams to search against the KOfam HMM database of Kyoto Encyclopedia of Genes and Genomes (KEGG) orthologs (58) and automatically assign hits above the KEGG bitscore thresholds for each KOfam profile. Additionally, we ran anvi-run-ncbi-cogs to search against the NCBI Clusters of Orthologous Groups (COGs) database (59) and identified archaeal single-copy core genes using anvi-run-hmms -I Archaea 76. To predict the presence or absence of metabolic pathways, we ran anvi-estimate-metabolism on each MAG. We annotated a metabolic pathway as present if over 70% of the genes in the pathway are present in a MAG, and partially present if 33–70% of the genes in a pathway are present in a MAG. Additionally, we searched for annotations of genes of interest within PROKKA annotations for each MAG, particularly for genes involved in fermentation, aerobic or anaerobic respiration, and energy metabolism. Sequences belonging to genes of interest were retrieved from each MAG and further inspected.

Protein sequences belonging to positive hits for denitrification genes from ODZ DPANN MAGs were obtained for *nosZ*. We extracted and aligned with MAFFT v7.450 using the --auto and --leavegappyregion parameters. Alignments were visualized in JalView 2.11.2.6 (60) and inspected for alignment quality and the conservation of key enzymatic regions for *nosZ*. Prediction of membrane-bound regions, protein localization, and protein structure were determined via DeepTMHMM (61). To create a protein tree for *nosZ*, bacterial and archaeal *nosZ*-encoded protein sequences were obtained from NCBI using the query esearch -db protein - query "*NosZ*" | efetch -format fasta, and sequences under 200 amino acids and over 800 amino acids were removed. In addition, cytochrome c oxidase subunit II proteins from bacteria and archaea were downloaded from NCBI using the queries esearch -db protein -query "cytochrome c oxidase subunit ii [PROT] AND bacteria [ORGN]" | efetch -format fasta and esearch -db protein -query "cytochrome c oxidase subunit ii [PROT] AND archaea [ORGN]" | efetch -format fasta. Sequences under 200 amino acids and over 800 amino acids were removed. To remove redundant or very similar sequences, Usearch v11 was used to cluster NCBI *nosZ* and Cox2 sequences at 90% amino acid identity with the flags -cluster_fast -id 0.9 -centroids (62). From clustered Cox2 sequences, 15 Cox2 sequences from taxonomically diverse organisms were chosen at random. These selected Cox2 were concatenated with clustered *nosZ* sequences and DPANN *nosZ* sequences and aligned with MAFFT v7.450 (63) using the --auto and -- leavegappyregion flags. The resulting Cox2 and *nosZ* alignment was trimmed with trimAl 1.4.1 with the -automated1 flag (64). We used the trimmed alignment to create a maximum likelihood protein tree using IQ-Tree v1.6.12 with 1000 ultrafast bootstraps.

### Methods for modeling producer and consumer dynamics

We used COMSOL (v5.6) to simulate the concentration field and associated uptake rate around a two-cell system consisting of a producer and consumer in three-dimensional space. In this simulation, the producer cell is represented as a sphere with a constant normalized concentration of 1 on its surface. The consumer cell on the other hand is represented by a sphere with a normalized concentration of 0 on its surface. We represent all aqueous concentrations as relative concentrations between producer and consumer cells to keep the system universal. We then strategically vary the relative radius of the producer (*R*) and consumer (*r*) cells, and the distance between the surface of the two cells (*d*) to disentangle the influence of these different factors on the relative substrate uptake of the consumer cell. A list of parameters and their values can be found in Table S1 which we cross-combine to create a total of 100 simulations. The whole simulation domain is a square domain of 20 µm side length with the consumer and producer cells equidistant from the center in the horizontal plane. All simulations are solved for steady-state concentration fields and the total uptake rate of the consumer cell is calculated directly in COMSOL.

### Genetic engineering methods and determining N_2_O concentrations

We inserted DNA sequences derived from DPANN putative nitrous oxide reductase genes into a *Pseudomonas aeruginosa* strain PA14 model system on a plasmid integrated into the genomic attTn7 site (65). Production and consumption of N_2_O by these cultures was quantified using a microelectrode (Unisense, Denmark). Details are in Supplementary Methods.

## Results

### DPANN within the ODZ archaeal community

From a set of 962 MAGs >50% completion and <10% contamination binned from the ETNP and Arabian Sea ODZs (46), 33 MAGs were taxonomically assigned to the DPANN superphylum, with 23 Woesarchaeota, 2 Pacearchaeota, 2 Nanoarchaeota, 1 Iainarchaeota, 3 Undinarchaeota, and 2 MAGs assigned to SpSt-1190, also known as *Candidatus* Altiarchaeota. The novel SpSt-1190 phylum was previously characterized in hydrothermal vents (10) but not in marine water columns. While our Woesearchaeota and Pacearchaeota MAGs were classified by GTDB-tk as members of Nanoarchaeota, phylogenetic analyses confirm their placement within these phyla (Figure 2). DPANN MAGs mapped to ODZ metagenomes within all three ODZs, including ETSP and ETNP metagenomes spanning multiple cruises, sampling sites, and years (Figure 1A). However, no DPANN MAGs were recovered from oxygenated surface metagenomes from the ETNP. Searching the TARA Oceans dataset comprising 957 non-redundant MAGs from co-assemblies from the global surface oceans (<10 m depth) and deep chlorophyll maxima (10–100 m depth) reveal no MAGs belonging to ODZ DPANN groups, and only 2 DPANN MAGs, both of which originated from the Red Sea and were assigned to Halobacteriota. The remainder of the 87 archaeal MAGs from TARA Oceans were assigned to either Thermoplasmatota or Thermoproteota. From 962 ODZ MAGs, 169 archaeal MAGs include 133 Thermoplasmatota or Thermoproteota, 2 Hydrothermarchaeota, and 1 Methanobacteriota to complement the 33 DPANN archaea. The average completion of retrieved ODZ DPANN MAGs was 75% with an average contamination of 2.6%. Dereplication at 99% average nucleotide identity (ANI) resulted in 16 unique MAGs (1 Iainarchaeota, 2 Nanoarchaeota, 1 SpSt-1190, 3 Undinarchaeota, 1 Pacearchaeota, and 8 Woesarchaeota).

**Figure 2:**
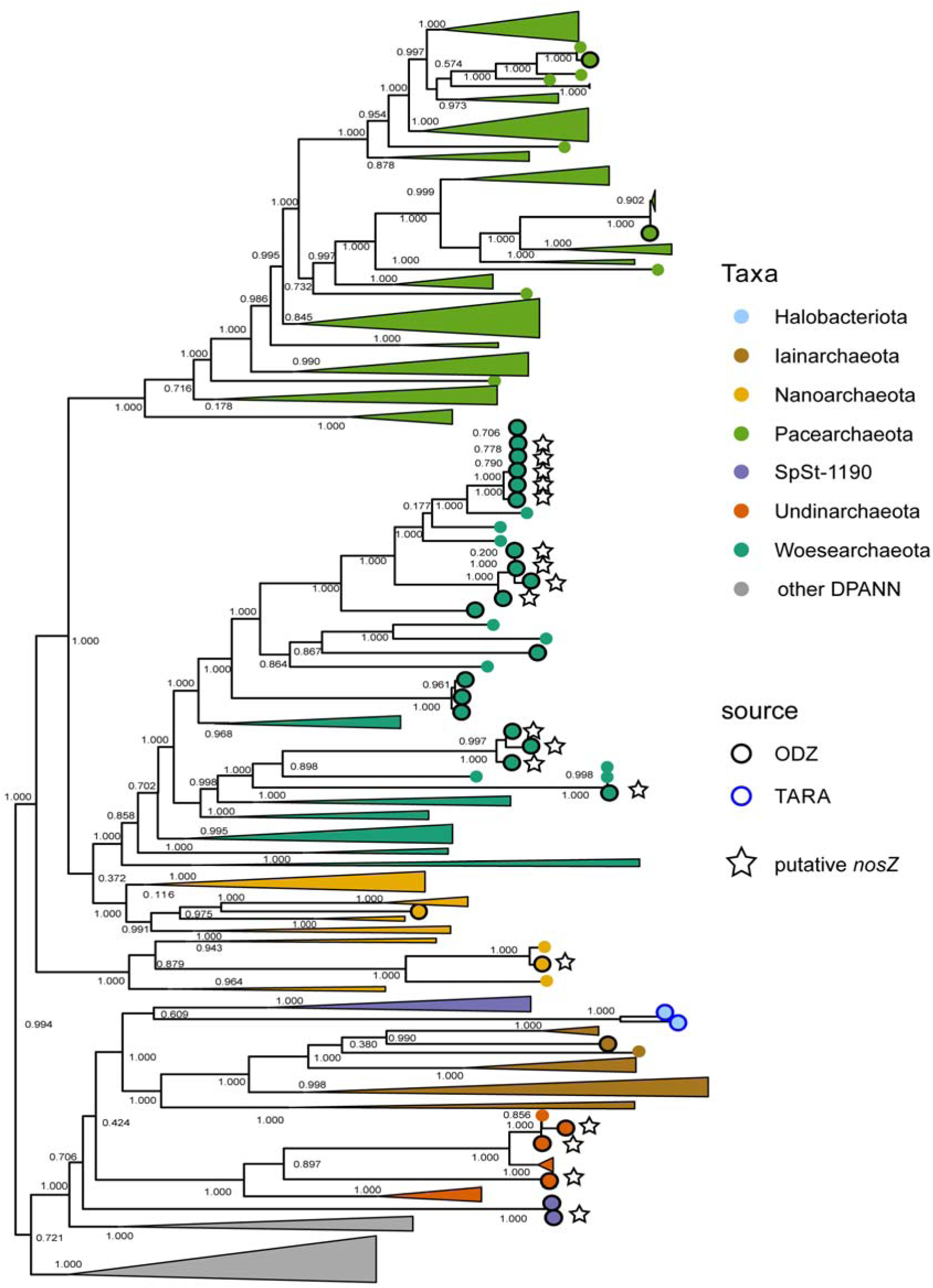
Species tree of DPANN MAGs and genomes from JGI, NCBI, ODZs, and TARA Oceans collections, colored by phylum-level taxonomy. Only DPANN phyla containing ODZ or TARA MAGs are shown. Black outlined circles indicate ODZ MAGs, blue outlined circles indicate TARA Oceans MAGs, and circles without outlines indicate NCBI or JGI genomes. Stars next to tips indicate the presence of a putative *nosZ*-like gene. Numbers by nodes correspond to bootstrap supports.

The archaeal population within ODZs, as represented by mapping to non-redundant archaeal ODZ MAGs, peaks within the oxycline above the ODZ core, with archaeal MAGs comprising 14% of the microbial community at 80 m depth in the ETNP (Figure S1A). At these depths, Thermoplasmatota and Thermoproteota dominate, with DPANN MAGs present at 0.25% relative abundance or less. DPANN MAGs also mapped to few reads from surface metagenomes from either the ETNP or ETSP, indicating low or absent populations of ODZ DPANN groups in surface waters (Figure 1B). No surface waters from the Arabian Sea were sampled. Within the anoxic ODZ core, DPANN archaea comprise about 25–50% of the archaeal community (Figure S1B). Highest relative abundances occur around 200 m depth in the ETNP (about 1% of the total community and 50% of the archaeal community) and the Arabian Sea (about 0.8% of the total community and 27% of the archaeal community) (Figures 1B, S1B). In the ETSP, relative abundances are lower (about 0.3% of the total community and 25% of the archaeal community), but peak approximately at the same depths (100–200 m). While abundances and distributions are similar across the various ODZs, the Arabian Sea harbors a comparatively larger proportion of Nanoarchaeota, although Woesarchaeota are still the most abundant fraction in general. The ETSP and ETNP are primarily dominated by Woesarchaeota with smaller contributions by Pacearchaeota, SpSt-1190, and Undinarchaeota (Figure 1B).

A phylogenetic tree of ODZ DPANN MAGs along with MAGs and genomes retrieved from NCBI and JGI reveals that ODZ Woesarchaeota MAGs fall within one primary clade, although several Woesarchaeota MAGs branch within other groups (Figure 2). Sister taxa falling next to ODZ Woesarchaeota derive from Mariana Trench surficial sediments, coral reefs, and groundwater metagenomes from NCBI. However, ODZ Pacearchaeota and Nanoarchaeota MAGs fall within distinct clades, indicating that these MAGs, despite belonging to the same phyla, are not closely related. The two SpSt-1190 MAGs from ODZs branch outside of the SpSt-1190 clade, and potentially form a distinct lineage from other SpSt-1190.

### Carbon metabolism within ODZ DPANN archaea

A metabolic analysis of ODZ DPANN MAGs shows diverse metabolic capabilities across MAGs, but limited metabolic and biosynthetic pathways within each MAG (Figure 3). Metabolic capabilities described are based upon annotations against the KEGG and COGs databases and require functional verification. While these annotations are predictions only, they offer estimates of metabolic potential for these uncultured organisms. Regarding anabolic synthesis, MAGs belonging to Nanoarchaeota have the most limited capabilities, with absent glycolysis of 3-carbon compounds, no TCA cycle genes, no pentose phosphate pathways, no pathways detected for the biosynthesis of most amino acids, and limited biosynthetic pathways for purine nucleotides. The absence of these pathways, even when considering these as partial genomes, suggests extremely limited abilities to synthesize purines, amino acids, lipids, vitamins, and other necessary cellular components. Other DPANN MAGs possess more metabolic capabilities, although most lack evidence of complete glycolysis, TCA cycle, and pathways for synthesis of multiple essential amino acids. Our draft Woesarchaeota genomes possess partial or complete capabilities for the last stages of glycolysis, the non-oxidative or reductive portions of the pentose phosphate pathway, and pyruvate oxidation. Additionally, most are capable of partial or complete purine and pyrimidine biosynthesis. Other central carbon archaeal pathways vary, with most MAGs lacking the shikimate pathway for biosynthesis of aromatic amino acids (66), the biosynthesis pathway for the ubiquitous cofactor coenzyme A, and the DeLey-Douderoff pathway for galactose utilization, which is analogous to the Entner-Doudoroff pathway (67). Several MAGs lack the ability to synthesize the intermediate phosphoribosyl diphosphate (PRPP) used in building nucleotides, some amino acids, and essential cofactors (68), as well as biosynthesis pathways for isoprenoids (Figure 3).

**Figure 3:**
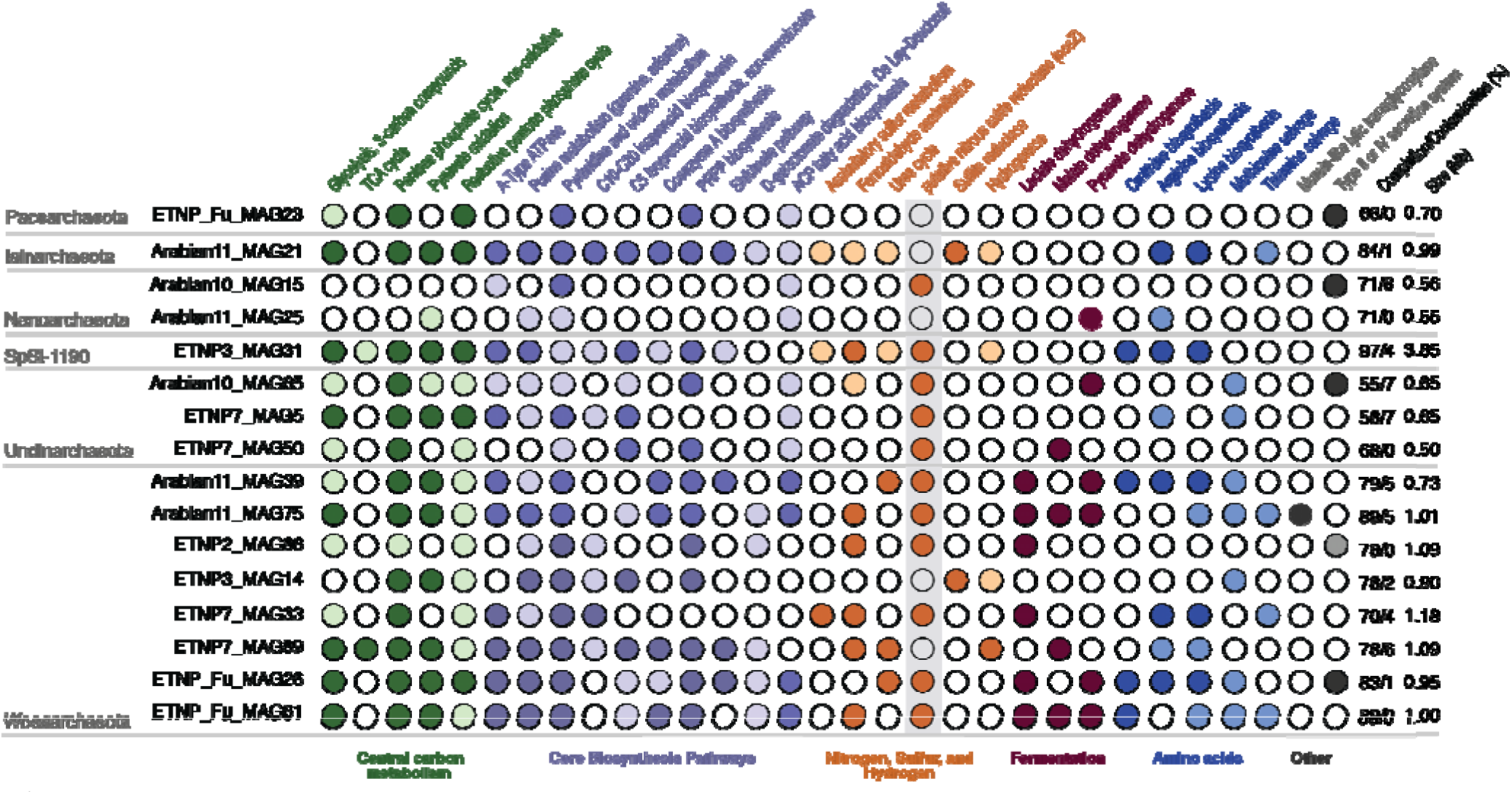
Metabolic analysis of unique DPANN MAGs. Circles show the presence/absence of key metabolic pathways, grouped by color according to general metabolism categories. Darker circles indicate >70% of genes within the pathway are present, while lighter circles indicate partial pathways (33–70% present). White circles indicate <33% of genes are present, and the pathway is considered absent. Completion/contamination and size of MAGs are shown on the right.

In accordance with other published DPANN archaea (4, 7, 8), the genome sizes of most ODZ DPANN are small, averaging 1.05 Mb. The exception are MAGs belonging to SpSt-1190, which have genome sizes of 4 Mb. DPANN MAGs encode for a number of transporters, including ones for zinc, iron, magnesium, and other metals, biotin transporters, SemiSWEET transporters for cellular uptake and translocation of sugars, and other ABC-type transporters.

Peptidases, particularly signal peptidases and membrane-bound peptidases, are also widespread. 7 DPANN MAGs from 4 phyla contain genes for Type II or IV secretion systems associated with protein transport and DNA exchange across membranes. Additionally, 3 DPANN MAGs encoded a murein-like lytic transglycosylase (1, 69). Normally absent in archaea, these large proteins bind and degrade peptidoglycan strands such as in bacterial cell walls (70).

Several DPANN MAGs possess the 3-oxoacyl-ACP reductase FabG, enoyl-ACP reductase FabI, and 3-hydroxyacyl-ACP dehydratase FabZ. These acyl carrier protein (ACP) fatty acid biosynthesis genes are typically found within bacteria and eukaryotes, which possess a bacterial pathway for lipid biosynthesis, the methylerythritol phosphate (MEP) pathway, while typical archaea use the non-homologous mevalonate (MVA) pathway. This “lipid divide” is a central distinguishing feature between archaea and bacteria (3). We find a distinction between Woesarchaeota and Nanoarchaeota MAGs, which possess the MEP pathway, while SpSt-1190, Undinarchaeota, Pacearchaeota, and Iainarchaeota contain genes for the MVA pathway. Further BLAST searching of DPANN ACP pathway proteins against the NCBI non-redundant protein database reveal high sequence similarity to other protein sequences from DPANN archaea, although the next-highest scoring hits primarily belong to bacteria. Additionally, 5 Woesarchaeota DPANN MAGs carry genes for cyclopropane fatty acid phospholipid synthesis, which are used in stabilizing bacterial phospholipid membranes (71) but have not been previously reported in archaea.

### Energy metabolism and nutrient cycling within ODZ DPANN archaea

Lactate, malate, and pyruvate dehydrogenases were present in 7 out of 8 unique Woesarchaeota and 2 out of 3 unique Undinarchaeota MAGs, indicating fermentative capabilities (Figure 3). These genes were absent in most Pacearchaeota, Nanoarchaeota, SpSt-1190, and Iainarchaeota MAGs. Two DPANN MAGs also carried the A and B subunits of assimilatory anaerobic sulfite reductase, but none carried dissimilatory sulfur cycling genes. Additionally, several DPANN MAGs contain desulfoferredoxin, manganese superoxide dismutases, and thioredoxin, which are involved in antioxidant systems (14), despite living in anoxic water columns. Formaldehyde assimilation genes were found within a number of Woesarchaeota MAGs, suggesting the ability to use one-carbon compounds for growth. MAGs belonging only to SpSt-1190 also encoded nearly-complete pathways for methanogenesis, along with a number of other methane metabolisms including the ribulose monophosphate pathway and methanofuran biosynthesis.

Several Woesarchaeota MAGs encoded hydrogenases, with 1 MAG encoding an FeFe-type hydrogenase potentially used in fermentative metabolism, while 2 encode an NiFe-type hydrogenase that may catalyze hydrogen oxidation for energy, which has recently been shown to be widespread among archaea (71) and marine bacteria (72). Additionally, several ODZ DPANN contain genes for urea cycling. These metabolic capabilities indicate diverse roles in carbon, sulfur, hydrogen, and nitrogen cycling for DPANN archaea within ODZs.

### Potential nitrous oxide reduction capability within ODZ DPANN

Within our 33 DPANN MAGs, 21 encoded a gene annotated as nitrous oxide reductase (*nosZ*), which catalyzes the reduction of N_2_O to N_2_. An HMM search against the HMM profile from validated *nosZ* sequences returns expectation values between 8.4 × 10^-17^ and 2.2 × 10^-5^ and bit scores from 57.9–20.1, compared to canonical *nosZ* e-values of less than 1.2 × 10^-33^ and bit scores > 113. In comparison, cytochrome c oxidase subunit II proteins returned expectation values of 0.003 – 7.5 × 10^-5^ and bit scores of 13–18.4. While bit score cutoffs vary, a bit score >50 is considered almost always significant (73).

However, other denitrification genes were absent within these MAGs. A gene tree built with canonical *nosZ* from bacteria and archaea, the DPANN *nosZ*-like protein, and the closely related homologue cytochrome c oxidase subunit II protein (Cox2) indicates DPANN *nosZ*-like genes comprise a monophyletic clade branching in between Cox2 and clade II Sec-type *nosZ* (Figure 4). Further investigation of multiple-sequence protein alignments of *nosZ*-encoded nitrous oxide reductase and the DPANN *nosZ*-like protein reveal the presence of a conserved copper-binding site, the Cu_A_ site, which has been reported within *nosZ* and Cox2 proteins. This site is exemplified by the C_1_X_3_C_2_X_3_H binding motif (74–76). The Cu_Z_ catalytic site typically found within *nosZ* is not found within the DPANN *nosZ*-like proteins, and DPANN nosZ-like proteins are shorter (56–617 amino acids) than canonical *nosZ* proteins (200–796 amino acids, although sequences vary in completeness). The Cu_Z_ site, which lacks a specific conserved motif, is characterized by 7 conserved histidine residues (35, 76, 77). Within 2 DPANN MAGs, we find a short cupredoxin-like domain protein directly upstream of the *nosZ*-like protein containing 5 conserved histidine residues, which clusters with clade II *nosZ* sequences containing the Cu_Z_ site within the protein phylogeny (Figure 4).

**Figure 4:**
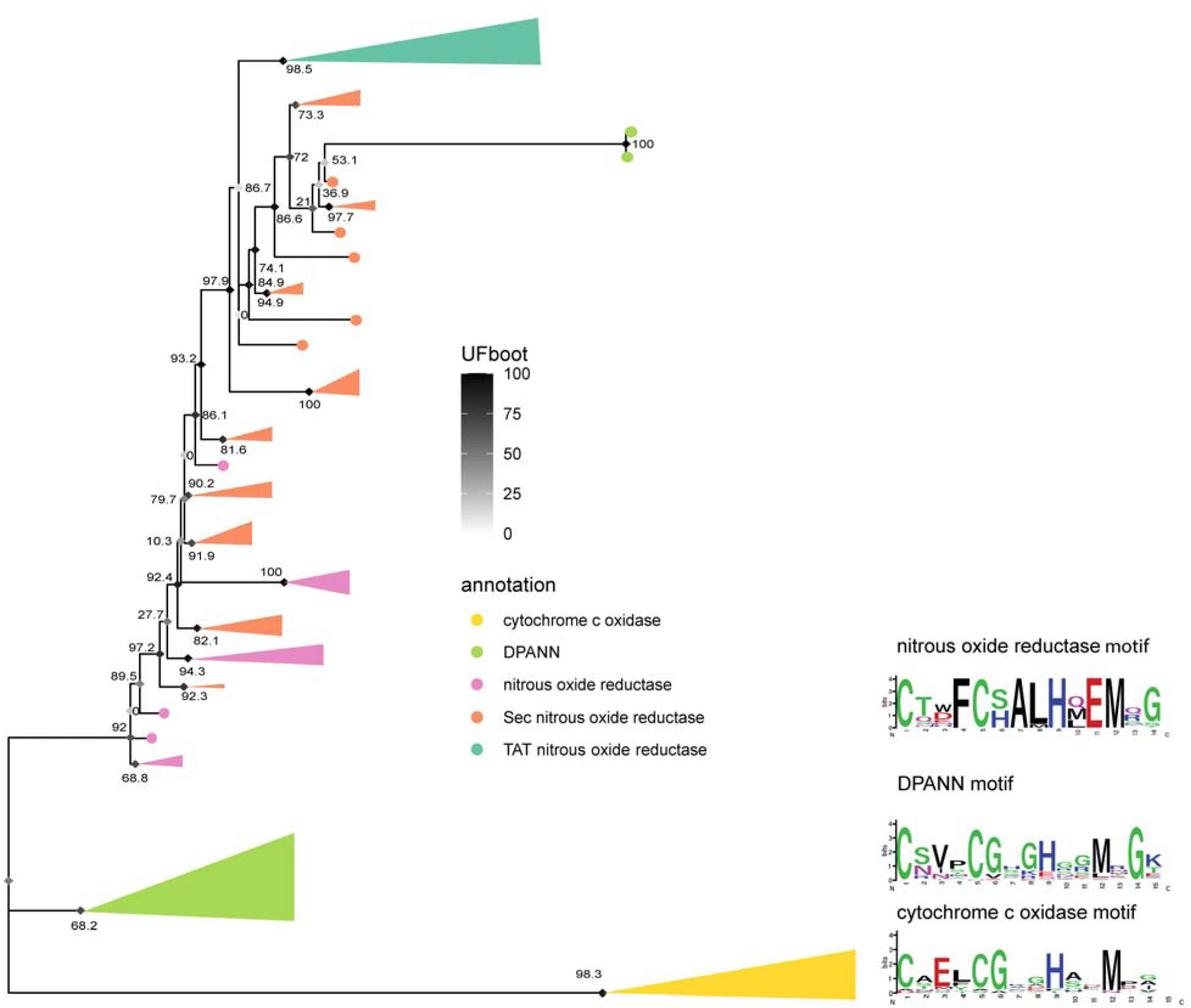
Protein tree of DPANN *nosZ*-like proteins (green) within the larger tree of canonical *nosZ* proteins (typical TAT type in teal, atypical Sec type in orange, type unknown in pink). Tree is rooted on cytochrome c oxidase subunit II (Cox2) proteins, shown in yellow. Diamonds at nodes correspond to ultrafast bootstrap supports, while numbers are SH-alrt values. Sequence motifs for the conserved Cu_A_ copper-binding site for each protein are displayed at right.

The mature NosZ protein resides in the periplasmic space in known denitrifiers (35). Predictions of protein location for NCBI *nosZ* sequences and the DPANN *nosZ*-like gene indicate both contain a signal sequence followed by the majority of the protein located outside the inner membrane, perhaps indicating function more similar to *nosZ* (Figure S2). In contrast, the Cox2 protein contains two transmembrane regions. A heterologous complementation test performed by separately introducing 3 DPANN *nosZ*-like sequences into a *Pseudomonas aeruginosa* Δ*nosZ* mutant did not yield significant differences in N_2_O consumption between *P. aeruginosa* Δ*nosZ* with DPANN *nosZ*-like gene insertion vs. the Δ*nosZ* parent strain (Figure S3). One strain carrying a putative DPANN *nosZ*-like gene variant displayed reduced N_2_O concentrations compared to the Δ*nosZ* parent, indicating potential N_2_O consumption, but this difference was not statistically significant (*p* = 0.26). Our positive control, the wild type *P. aeruginosa* PA14 containing a functional *nosZ* gene, displayed reduced N_2_O concentrations (*p* < 0.001), validating the ability of our experimental system to detect N_2_O accumulation.

N_2_O in the ODZ typically exists at bulk nanomolar concentrations (78), raising the question of whether specialization on N_2_O consumption is metabolically feasible. However, local N_2_O concentrations may vary in the presence of N_2_O producers such as partial denitrifiers carrying upstream denitrification capabilities. We simulate the conditions under which local N_2_O concentrations differ from bulk conditions by varying a set of parameters representing the distance between the N_2_O consumer and N_2_O producer and the size ratio of the two cells (Figure 5). Generally, two conditions favor elevated N_2_O uptake rates for the consumer normalized to cell volume: when the consumer cell is small relative to the producer cell (Figure 5B) and when the distance between the producer and consumer cells is small (Figure 5C). Rates are normalized to cell volume to reflect the important consideration that resource requirements are proportional to cell size (79). Consumer cell size has a decisive influence on N_2_O uptake rate as a smaller cell size both increases the uptake rate normalized to cell volume and a large producer-to-consumer size ratio surrounds the consumer cell within the diffusive boundary layer of the producer cell. For example, an increase in the ratio of consumer-to-producer radii from ∼ 0.1 to ∼1 results in an average 100-fold decrease in N_2_O uptake for attached consumer cells. Similarly, increasing the producer cell size three-fold increases the attached consumer N_2_O uptake rate between ∼14% for small consumer cells to 67% for larger consumer cells. Incredibly, increasing distances between cells from 0 µm to 0.1 µm reduces the maximal N_2_O uptake rate by ∼65% on average, and a consumer cell merely 2 µm from a producer receives 93% less than those attached (Figure 5C). Therefore, when bulk N_2_O is low as in the ODZ (80), consumer cells experience high N_2_O supply only when they are in physical contact with an N_2_O producer (*d* = 0 µm) and when they are much smaller in size relative to the producer, as would be the case for episymbiotic DPANN archaea.

**Figure 5:**
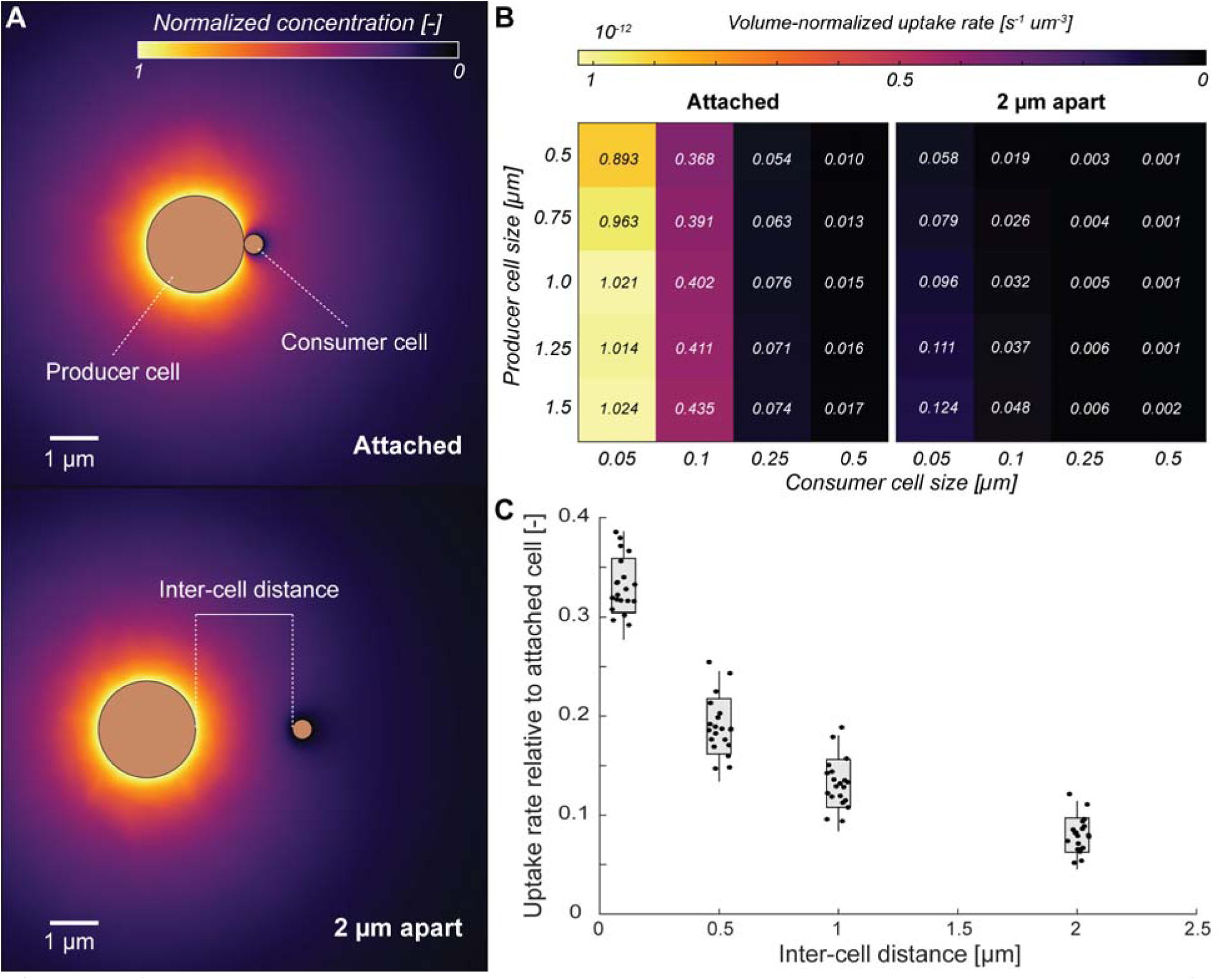
**A)** Schematic showing the spatial N_2_O concentration for two inter-cell distances of *d* = 0 µm (attached) and *d* = 2 µm (free-living). The relative surface N_2_O concentration for the producer is set to 1, while the relative surface N_2_O concentration of the consumer is set to 0. The radius of the producer, radius of the consumer, and distance between the cells are varied according to values in Table S1. **B)** Volume-normalized uptake rate of N_2_O for the consumer at 0 µm separation (attached) and 2 µm separation (free-living) for all values of the consumer and producer cell sizes. Numbers indicate the actual volume-normalized uptake rates (multiplied by 10^-12^). **C**) Uptake rates as a function of the inter-cell distance normalized to the attached scenario of the same consumer-producer cell size combination. A value of, e.g., 0.2 indicates that this combination of producer and consumer cell size shows a reduction of 80% in the consumer N_2_O uptake rate at this distance compared to if they were attached. n = 20 simulations plotted for each bar, with box representing ± 1 s.d. and the whiskers showing ± 2 s.d.

## Discussion

DPANN archaea were found to be a stable resident population within all 3 permanent pelagic ODZs. Abundances of DPANN archaea, including Nanoarchaeota, SpSt-1190, Iainarchaeota, Woesarchaeota, and Undinarchaeota, increase as oxygen decreases, while few or no DPANN archaea were found in the surface oceans (Figure 1B). While a few population differences are found between ODZs, Woesarchaeota are the dominant phylum within all three ODZs, with Nanoarchaeota in the Arabian Sea and Pacearchaeota, Undinarchaeota, and SpSt-1190 in the ETNP and ETSP forming the second most abundant groups (Figure 1B).

ODZ DPANN archaea are phylogenetically and metabolically diverse and group together with other DPANN from non-ODZ environments, although several Woesarchaeota cluster within the same clade (Figure 2). Similar to DPANN across various environments (4, 10, 12, 13), most ODZ DPANN have small genome sizes and limited capacity for biosynthesis of essential amino acids and nucleotides, limited energetic capabilities, and partial or absent pentose phosphate pathways despite overall high MAG completion estimates (Figure 3). Additionally, completion metrics may underestimate the completeness of DPANN MAGs due to their limited genomes and high number of absent genes considered essential in other organisms. Numerous studies have reported microscopy images of environmental DPANN attached to host cells (7, 13, 81). While most ODZ DPANN genomes suggest a host-associated rather than free-living lifestyle, SpSt-1190 genomes average 4 Mb in size, possess a number of biosynthesis pathways, and carry pathways for methanogenesis. These unique archaea may represent free-living DPANN organisms (82–84) involved in methane cycling. The identification of ODZ DPANN hosts, whether a single host, various hosts, or a community, may hold keys to their distribution and survival within ODZ environments.

Studies have suggested a role of DPANN archaea for carbon cycling, such as by scavenging organic carbon in the form of nucleotides, lipids, and amino acids (21, 85), participating in the exchange of carbon compounds with hosts, and even directly parasitizing upon hosts (12). In addition, some may perform fermentation and consume or produce acetate

(6). We find conserved pathways for amino acid salvage and fermentation across ODZ DPANN genomes (Figure 3). While various sugar, protein, and DNA transporters indicate potential resource exchange with host cells, the existence of a peptidoglycan-degrading enzyme and secreted peptidases within several MAGs may point to a potentially parasitic relationship between host and DPANN cell. Future experimental tests will be needed to clarify these metagenomic predictions.

While other nitrogen cycling genes are absent, a majority of ODZ DPANN carry a gene similar to the nitrous oxide reductase gene *nosZ* that catalyzes the reduction of nitrous oxide (N_2_O) to N_2_. Further investigation of this gene, annotated as nitrous oxide reductase, indicates the presence of a conserved Cu_A_ copper-binding site typical of *nosZ* and cytochrome c oxidase subunit II (75, 76) (Figure 4). The cellular location of the protein product of the DPANN *nosZ*-like gene is postulated as outside of the membrane, possibly in the periplasmic space (Figure S2). DPANN archaea are thought to possess two membranes (86), and canonical *nosZ* is a periplasmic protein unlike the membrane-bound cytochrome c oxidase subunit II (74). Cytochrome c oxidase performs the last step of aerobic respiration, but no other elements of aerobic respiration were found within these archaea (Figure 3). The Cu_Z_ catalytic center, typically found upstream of the Cu_A_ center in *nosZ*, is absent within DPANN *nosZ*-like genes. The Cu_Z_ center lacks a consensus motif, but is characterized by 7 histidine residues that bind copper ions (87). While the majority of DPANN MAGs possess several acyl carrier protein genes for fatty acid biosynthesis surrounding the *nosZ*-like gene, 2 DPANN MAGs encode a protein containing 5 histidine residues directly upstream of the *nosZ*-like gene. This protein, annotated to the same family as *nosZ*, groups phylogenetically with clade II *nosZ* sequences (Figure 4) and may perform a function related to that of the Cu_Z_ site. This hypothetical histidine-rich region was absent within other DPANN MAGs, and was not included within the complementation test. The activity of these or other proteins within these genomes may be required for N_2_O reduction. While the function of putative *nosZ*-like genes within DPANN archaea remain hypothetical, evidence suggests the involvement of these genes in N_2_O reduction or another redox process with metabolic or physiological importance due to their conservation within these small, streamlined genomes.

Complementation of *P. aeruginosa* Δ*nosZ* with DPANN *nosZ*-like genes did not result in significant N_2_O consumption. While heterologous complementation may offer convincing evidence for the function of unknown genes, negative results are difficult to interpret. Large evolutionary distances between DPANN archaea and the gram-negative bacterium *P. aeruginosa*, likely resulting in different intracellular conditions, may inhibit the proper transcription, translation, or maturation of the DPANN NosZ-like protein. The protein may also be adapted to specific environmental conditions necessary for its activity, which differ from those used during standard cultivation of *P. aeruginosa*. Deletion and complementation of the *nosZ*-like gene within native DPANN archaea would be an ideal functional test, but currently no cultured representatives or genetic toolkits are available for these organisms, limiting our knowledge of many of their metabolic features to predictions from gene annotations.

N_2_O exists in nanomolar concentrations in ODZs compared to the higher concentrations of nitrate and nitrite (78), posing challenges for N_2_O-reducing specialists lacking upstream denitrification genes. However, an N_2_O-consuming lifestyle may be feasible if local N_2_O concentrations are elevated in proximity to an N_2_O source, such as a partial denitrifier lacking *nosZ*. Previous studies have indicated widespread occurrence of partial denitrifiers lacking *nosZ* within ODZ regions (39, 46). We tested this scenario by modeling the local flux of N_2_O from a producer (the source) to an N_2_O consumer (Figure 5). N_2_O uptake rate of the consumer is elevated 100-fold when the two cells are in physical contact vs. when they are a short distance of 2 µm away (Figure 5C). This increase in N_2_O uptake rate drops off steeply, however, as the consumer-to-producer cell size ratio increases (Figure 5). Under low bulk N_2_O concentrations, partial denitrifiers may provide elevated local N_2_O only to much smaller surface-attached episymbiotic N_2_O consumers. DPANN archaea within ODZs, similar to those found within other environments (7, 8, 13), potentially exist as host-associated episymbionts and likely possess small cell sizes. The average cell volume of DPANN archaea has been reported as 0.004 µm^3^ (13) while average marine bacterial cell volume has been reported at up to 0.096 μm^3^ (88), resulting in a consumer-to-producer cell size ratio of < 0.05. Thus, DPANN archaea may be uniquely adapted to consume N_2_O and other resources that are scarce under bulk conditions but locally elevated in proximity to host cells.

DPANN archaea possess a high number of unknown or unannotated genes, representing “microbial dark matter.” Within our ODZ DPANN, we found over 20,000 hypothetical proteins across all MAGs. Further studies, possibly using genetic manipulations, isolation or enrichment cultures, imaging, and computational proteomics approaches are required to characterize the functions of putative or hypothetical proteins. The expanding knowledge of these organisms may make these questions more tractable in the near future. At a large scale, the scavenging of carbon, potential nitrogen, sulfur, and hydrogen cycling capabilities, and ecological effects on host populations via symbiosis or parasitism by DPANN archaea in the ODZs warrants future investigation.

## Acknowledgements

We thank Dr. Xin Sun (Carnegie Institution for Science) and Dr. Bess B. Ward, Dr. Amal Jayakumar, and Dr. Samantha G. Fortin (Princeton University) for sample collection, DNA extractions, and providing the resulting metagenomics data we used to assemble MAGs for this study. Funding for this project came from Simons Foundation award 622065 and National Science Foundation award OCE-2142998 to ARB. IZ was supported in part by an MIT School of Science MathWorks Science Fellowship. Grants to SW (from the National Science Foundation Graduate Fellowship Program) and to DKN (from the NIH, R01 HL152190-03) also contributed. We are additionally grateful for the generosity of Dr. Bruce Heflinger in supporting the bablab, including this work.

## Author contributions

IHZ and ARB conceptualized this study. IHZ assembled metagenomes and MAGs, conducted bioinformatics analyses, and drafted the paper. BB and ARB conceived and carried out analyses regarding the N_2_O uptake model. RZ provided bioinformatics and overall guidance. DKN and SW conceived the heterologous complementation test for the *nosZ* homologs and provided all strains used in this study, and SW performed genetic engineering within the *Pseudomonas* model system.

## Supplementary Material

## Supplementary Methods

### Sequence selection and genetic engineering

DPANN *nosZ*-like genes from ODZ metagenome-assembled genomes were extracted and aligned with MAFFT-linsi v7.450 using the --leavegappyregion parameters (1). Three representative sequences were obtained by clustering the DPANN *nos*-like genes at 90% nucleotide identity with vsearch (2). Putative *nosZ*-like gene sequences were optimized for the *Pseudomonas aeruginosa* genome using the IDT codon optimization tool. These fragments were synthesized as gBlocks from Twist Bioscience. Using gBlocks as template, the fragments were amplified with homologous overhangs using PCR primers (Table 4.S1). Products were cloned into the pJM220 plasmid linearized by KpnI and HindIII digestion using Gibson assembly (NEB). Putative *nosZ*-like gene containing plasmids were introduced at the attTn7 site in the PA14(Δ*nosZ*) strain genome (3, 4). Strains with integrated plasmids will drive gene expression using a rhamnose-inducible promotor when grown in the presence of l-rhamnose.

### Growth conditions and N_2_O reduction tests

Three *Pseudomonas aeruginosa* PA14 Δ*nosZ* strains containing the putative *nosZ*-like genes were grown on LB plates at 37°C overnight along with the parent wild-type PA14 (positive control) and PA14 Δ*nosZ* (negative control). Colonies were inoculated into 10 mL Luria-Bertani (LB) broth within serum vials. Serum vials containing each strain, along with a cell-free control, were capped and sealed post-inoculation and grown within a 37°C incubator with shaking at 100 rpm. A total of 4 replicates were performed for each strain. After cultures used up the available oxygen to reach stationary growth phase, sterile 50 µM nitrate and 0.02% l-rhamnose w/v were injected into each serum bottle to drive denitrification and the expression of the *nosZ*-like genes. After 24 hours of growth at 37°C, N_2_O concentrations were measured using a Unisense N_2_O microelectrode connected to the Unisense Field Multimeter according to manufacturer protocols. N_2_O concentrations were normalized to the average N_2_O readout of the PA14 Δ*nosZ* control.

## Supplementary Tables

**Table S1:**
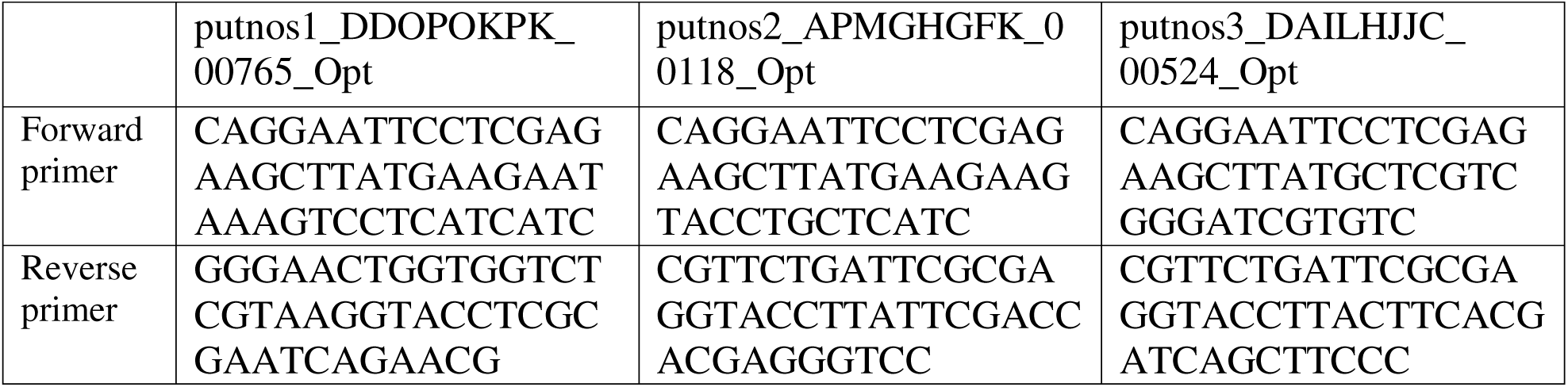
**Primers used to amplify DPANN *nosZ*-like gene fragments with overhangs for each of the 3 representative gene sequences.**

**Table S2:** **List of parameters used in model including specific values used for all simulations.**

| Parameter | Values |
| --- | --- |
| Producer radius R [$\mu\text{m}$] | [0.5, 0.75, 1.0, 1.25, 1.5] |
| Consumer radius r [$\mu\text{m}$] | [0.05, 0.1, 0.25, 0.5] |
| Cell distance d [$\mu\text{m}$] | [0, 0.1, 0.5, 1, 2] |

## Supplementary Figures

**Figure S1:**
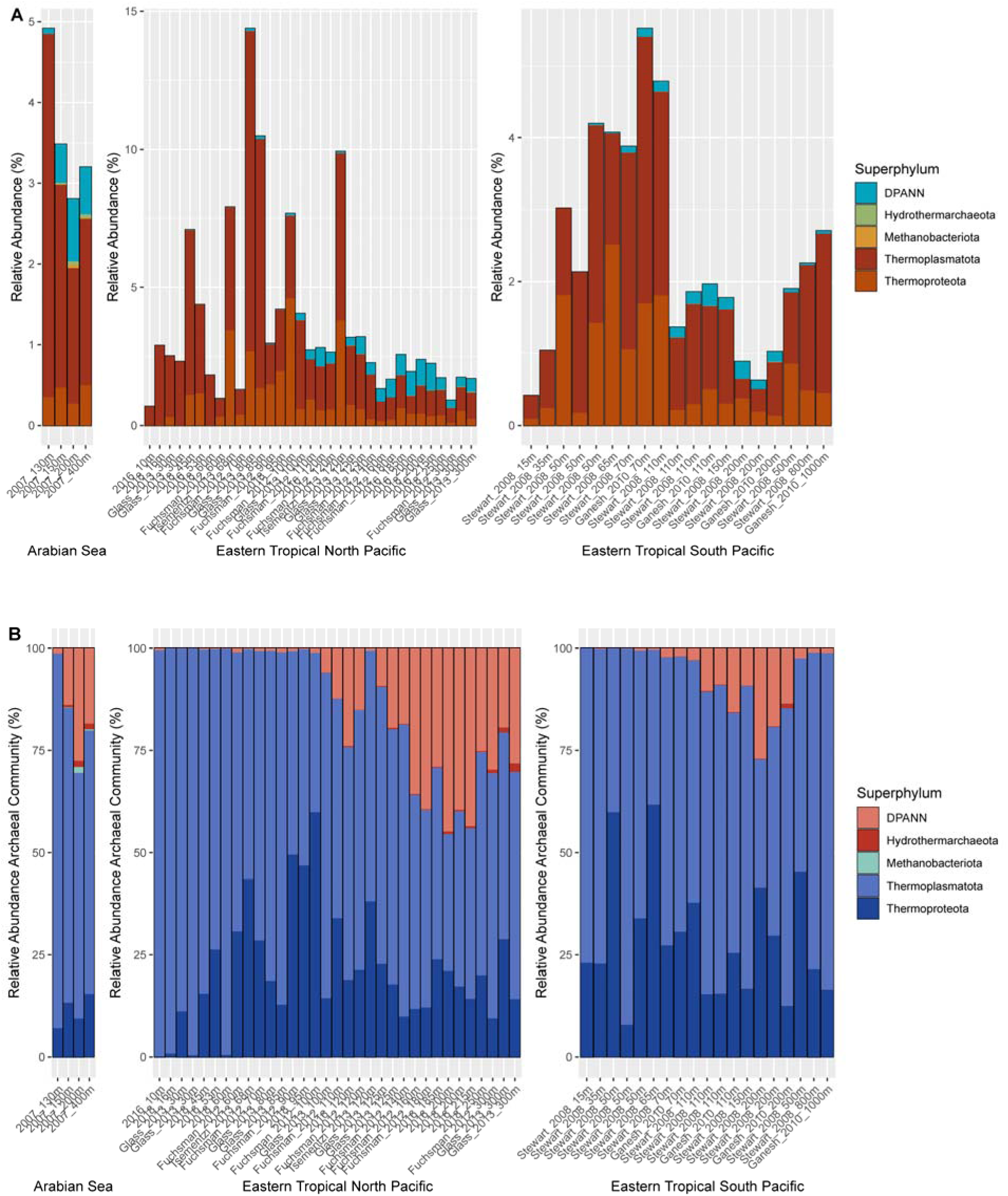
**A)** Percentage of the total community assigned to the Archaea domain, with DPANN archaeal phyla grouped together (teal). Other archaea are colored by phylum. **B)** Proportion of total archaeal reads belonging to the DPANN superphylum vs. other archaeal taxa, with DPANN archaeal phyla grouped together (orange). Other archaea are colored by phylum. All bars are scaled to 100%.

**Figure S2:**
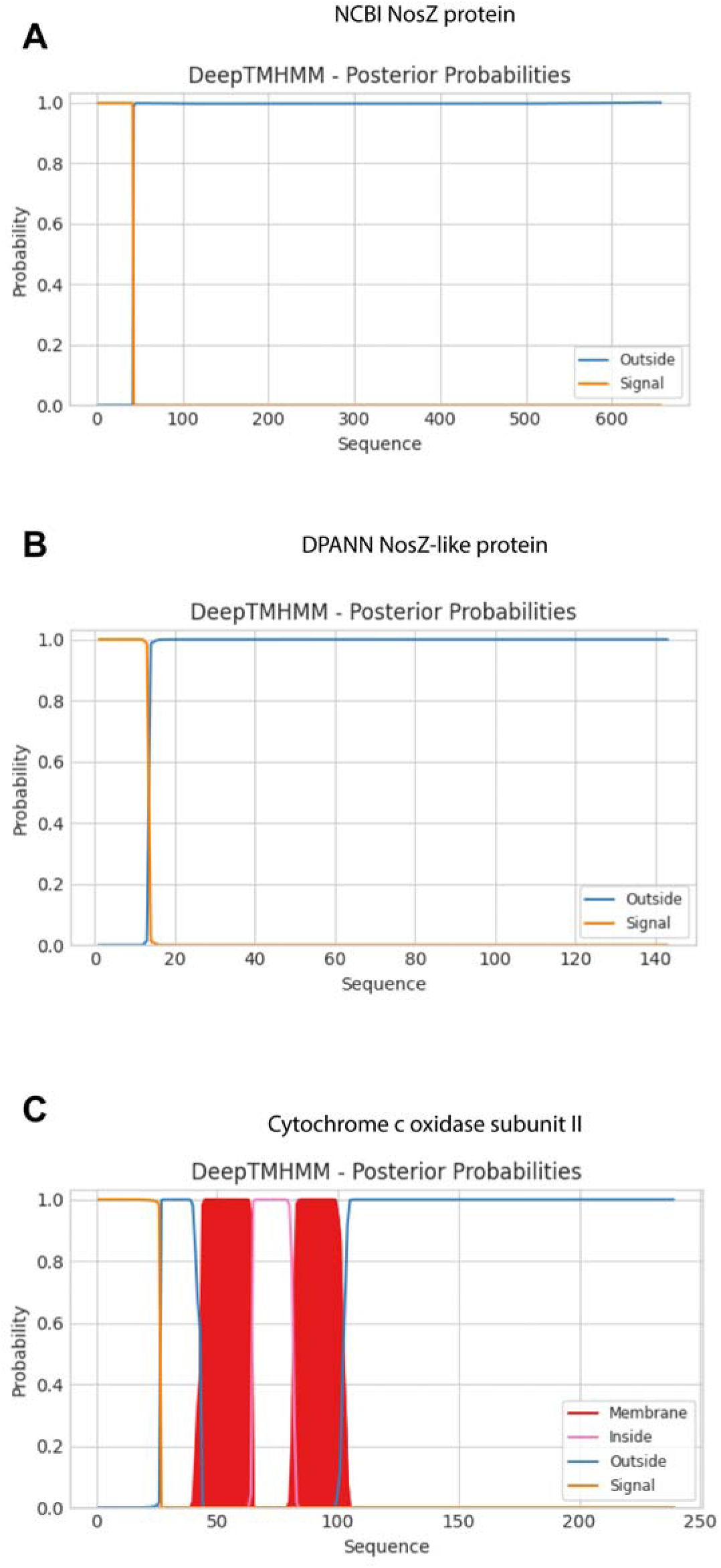
Prediction of protein location for **A)** canonical *nosZ*-encoded protein **B)** DPANN *nosZ*-like protein **C)** cytochrome c oxidase subunit II. Protein topologies and locations were predicted using DeepTHHMM. Probability scores on the y-axis indicate predicted probability of the protein region as belonging to a signal peptide sequence (orange), cytoplasmic (inside membrane, pink), membrane-associated (red), or outside the membrane such as in the periplasmic space (blue). Amino acid positions are indicated on the x-axis.

**Figure S3:**
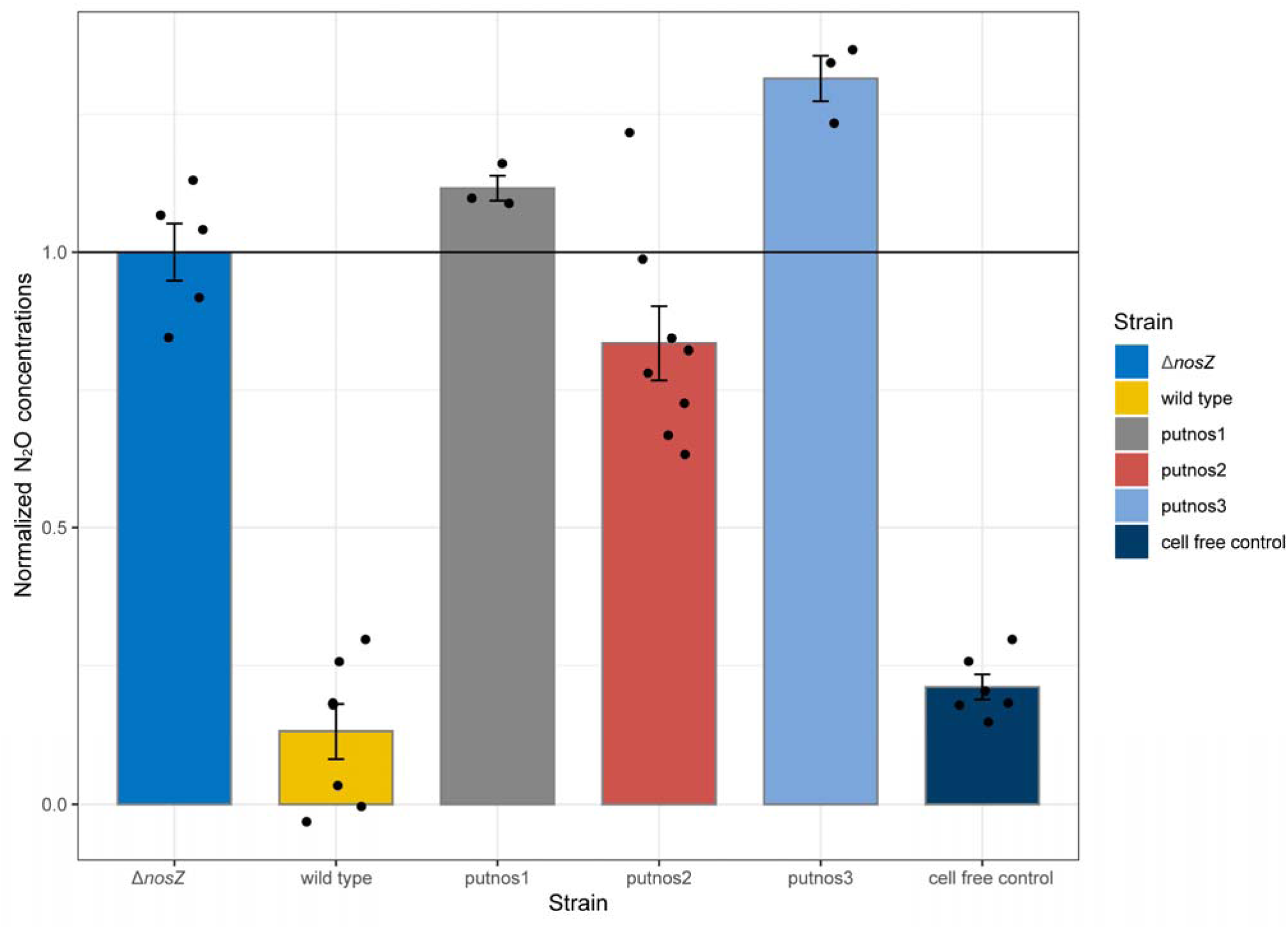
N_2_O concentrations after 24 hours of anaerobic growth in LB supplemented with 50 µM nitrate and 0.02% rhamnose to drive expression from the DPANN *nosZ*-like gene within the *P. aeruginosa* PA14 Δ*nosZ* background. N_2_O concentrations are normalized to the average N_2_O concentration for the parent PA14 Δ*nosZ* control. The putnos1, putnos2, and putnos3 strains correspond to the 3 representative *nosZ*-like genes after clustering DPANN *nosZ*-like sequences at 90% nucleotide identity. Wild-type PA14 and a cell free control are also shown.

